# Fast eco-evolutionary changes in bacterial genomes after anthropogenic perturbation

**DOI:** 10.1101/2020.03.13.990432

**Authors:** Manuel García-Ulloa, Ana Elena Escalante, Alejandra Moreno Letelier, Luis Enrique Eguiarte, Valeria Souza

## Abstract

Anthropogenic perturbations such as water overexploitation introduce novel selective pressures to the natural environments, impacting on the genomic variability of organisms and thus altering the evolutionary trajectory of its populations. Bad agricultural practices and defective policies in Cuatro Cienegas, Coahuila, Mexico, have strongly impacted its water reservoir, pushing entire hydrological systems to the brink of extinction together with their native populations. Here, we studied the effects of continuous water overexploitation on an environmental aquatic lineage of *Pseudomonas otitidis*, inhabitant to a particularly affected lagoon of an exhaustively studied system in the middle of the desert, over a 13 year period which encompasses three desiccation events. By comparing the genomes of a population sample from 2003 (original state) and 2015 (perturbed state), we analyzed the demographic history and evolutionary response of this bacterial lineage to the perturbation. Through coalescent simulations, we obtained a demographic model of contraction-expansion-contraction which, alongside an increment in mean Tajima’s *D* and recombination rate, loss of genetic and nucleotidic variation and a single amino acid under positive selection, points the occurrence of an evolutionary rescue event, possibly potentiated by horizontal gene transfer, where the population nearly went extinct during the first desiccation event but sharply recovered in the second and adapted to its new environment. Furthermore, the gain of phosphorylation, DNA recombination and small-molecule metabolism and loss of biosynthetic and regulation genes on the exclusive accessory genome suggest a functional shift to a more generalist scavenger lifestyle in an environment that went from oligotrophic to nutrient-rich.

## Introduction

Human population growth and human activities have negatively impacted fresh-water bodies globally [1, 2]. It is estimated that the water demand for human activities exceeds climate change as the most important factor influencing the state of water systems [3]. Among the different human uses for freshwater, agriculture is responsible for 70% of the global water extraction, a figure that may increase to 90% by 2050 due to drought and warmer temperatures associated with climate change [4, 5]. This negative feedback between diminishing fresh-water stocks, deforestation and climate change, has already caused droughts and desiccation events worldwide which in turn generate fragmentation and loss of habitat, putting the survival of many wild species at stake [6–8]

These extreme events caused by anthropogenic perturbations introduce novel and strong selective pressures; the response to these pressures depends on the standing genetic variation within populations [9]. In many cases, the lack of capability to adapt to the new environments impacts entire communities and ecosystems [10]. This “domino” effect is the reflection of the tight relationships and interactions among the different members of natural communities [10].

In bacteria, the best studied evolutionary response to an anthropogenic selective pressure is the rapid evolution of antibiotic resistance by multiple lineages of bacteria [11, 12], which has become a critical health issue worldwide [13]. Such fast adaptive response to new challenges in bacteria can happen along with SOS repair response leading to high mutation rates and abrupt declines in effective population sizes, involving genetic drift, fixating not only of new mutations, but also of new gene combinations acquired by horizontal gene transfer (HGT) [14, 15]. Although these processes may lead to the extinction of many bacterial lineages, they can generate new genetic combinations by changing the copy number of previously beneficial, neutral or even deleterious alleles in a process known as evolutionary rescue [16–18]. Also, evolutionary rescue can result in a selective sweep favoring the rapid expansion of a successful clone [19].

Arid ecosystems are particularly prone to deep water overexploitation by agriculture [20–21]. A dramatic example of this global trend is Cuatro Cienegas Basin (CCB), an oasis in the Chihuahua desert in northern Mexico. This system is composed of various hydrological systems that used to include numerous pools, lagoons and rivers fed by underground springs [22]. Nowadays, there is only 5% or less of the original wetland left, as the exploitation of water through canals that have drained the wetlands since the 70’s has had a strong impact on CCB hydrological systems. As an example of this tragedy, the larger lagoon of our long-term study site, Churince, no longer has water [23–25]. This is particularly poignant, since Churince was described as a “lost world” [24], where unusually unbalanced stoichiometry [26] had apparently isolated this wetland from modern microbial communities preserving a hyper diverse indigenous microbial community [24,27,28] of unique bacterial lineages of which a large fraction apparently descended from marine lineages [29–30]. We believe that these unique lineages survived and evolved by eco-evolutionary feedbacks [31] along with their complex communities that formed either microbial mats, stromatolites or, in the case of water, biofilms.

Churince system included three sequentially interconnected bodies of water: Laguna Grande, Laguna Intermedia and a spring. Laguna Grande, the most distal lagoon, dried up completely in 2006 [30]. Metagenomic studies performed in October 2012, showed the impact of desiccation in microbial communities of Laguna Intermedia: a decrease in biodiversity under dry conditions came along with an increase of negative interactions, mostly competition, among its members [25]. Likewise, changes in water stoichiometry, which may occur due to the increase in nutrients caused by the mortality of macro-organisms could have changed the dynamics of this normally oligotrophic site. An *in situ* nutrient enrichment experiment was performed in Churince using different proportions of N:P, showing that lower N:P ratio favored the proliferation of generalist bacteria and algae at the expense of the typically rare endemic lineages [32–34]. This was the case of pseudomonads, which appeared to benefit from the environmental disturbance [35, 36].

The genus *Pseudomonas* is characterized by having large genomes, high metabolic versatility and flexible gene expression thanks to its high percentage of regulatory genes, reaching up to 9.4% in *P. aeruginosa* [37]. This allows *Pseudomonas* to proliferate in a wide variety of environments, as well as plant and animal hosts including humans. Members of the genus, such as *P. aeruginosa* and *P. otitidis* [38], have been isolated from disturbed environments such as soils contaminated with hydrocarbons [39, 40], treatment plants of the dyeing industry [41], as well as septic tanks [42]. Therefore, their presence in high abundance is usually an indicator of the existence of an underlying environmental disturbance.

Biology and ecology of pseudomonads from Churince has been described in different studies [35, 43–45], and more recently two new strains of the *P. aeruginosa* PA14 clade native to Laguna Intermedia were sampled during the first desiccation event in 2011 [36]. *P. otitidis* from CCB represented a separate native lineage that was present, but was rare, before the perturbations at Laguna Intermedia [44].

In this study we took advantage of a unique opportunity to study the particular adaptations of pseudomonads in order to survive in the extremely oligotrophic environment of Laguna Intermedia during desiccation events, and to try to understand why this group of bacteria disappeared from the lagoon after October 2015 (even if the Laguna Intermedia did not completely dried until 2017). We used comparative genomics that allowed to reconstruct the evolution of *P. otitidis* at Laguna Intermedia in a period of 12 years, from the first sampling (2003) to the last time it was found (2015). In this time period Laguna Intermedia suffered four desiccation events that ended in 2017 when it finally dried up completely (and it remains so until today). We observed changes involving the pan-genome size, as the core genome increased and the genetic diversity decreased. SOS repair and an increase in recombination rate are likely involved in the observed changes. We used a coalescent analysis to understand the evolutionary changes in demography after the evolutionary rescue till the sudden final crash of the populations.

## Materials and Methods

### Study site

Laguna Intermedia from the Churince system in Cuatro Ciénegas, Coahuila, Mexico (26.848572 N 102.141783 W) suffered four recent desiccation events: October 2011, November 2012, October 2015 and October 2016 before completely drying up in 2017 [24, 36]. Churince was a long-term study site, and water physico-chemical conditions were recorded periodically since 2002 with the multiparameter water quality sonde Hydrolab MS5, until water disappeared from the two main lagoons in the system. In Laguna Intermedia, salinity, specific conductivity and pH were particularly altered with each desiccation event (S1 Fig).

### Sampling and isolate collection

On October 2015, samples were taken from surface water from multiple points of Laguna Intermedia using sterile BD Falcon vials (BD Biosciences, San Jose, CA). Isolates were obtained by plating 200 μl of each sample on *Pseudomonas* Isolation Agar (PIA) plates, which were further incubated at 32°C for 24 h. Individual colonies were transferred to new PIA plates and grown at 32°C for 1 day. Each colony was grown in LB medium at 32°C for 1 day and then stored in 50% glycerol solution at −80°C. From October 2015 until 2017, when it became completely dry, no other pseudomonad was isolated from Laguna Intermedia, even if they were intensively looked after in different subsequent samplings and cultures.

### Isolate identification

DNA extraction was done with Qiagen DNeasy Blood & Tissue kit and the 16S rRNA gene was amplified with universal primers 27F and 1492R. Seven colonies of October 2015 were identified as *P. otitidis* through a blast search against the RefSeq database. Additionally, five isolates from August 2003 previously identified as *P. otitidis* [45] were used for the analysis as representatives of the undisturbed state of the lagoon.

### Sequencing, assembly and genome annotation

Sequencing was carried out using the Illumina platform in its PE 2×300 modality, obtaining a coverage larger than 30x for all genomes. Genomes were assembled de novo with MaSuRCA 3.2.4 [46], followed by SPAdes 3.10.0 [47]. Contigs less than 600 bp long and 10x of k-metric coverage were discarded. The validation of the assemblies was done with Quast v4 [48] and BUSCO v2 [49]. All genomes were deposited on NCBI under the accession numbers JAANPJ000000000 (41M), JAANPK000000000 (39M), JAANPL000000000 (38M), JAANPM000000000 (36M), JAANPN000000000 (35M), JAANPO000000000 (34M), JAANPP000000000 (32M), JAANPQ000000000 (15M), JAANR000000000 (13M), JAANPS000000000 (12M), JAANPT000000000 (10M), JAANPU000000000 (9M). A database with all the complete annotated genomes of pseudomonas to date was built as reference for annotation with Prokka 1.11 [50].

### Genealogy of core genomes

Phylogenomic relationships between all lineages were assessed through a core genome genealogy. A genome alignment was done with progressiveMauve and the core genome was extracted with the stripSubsetsLCBs script [51]. Core genome genealogy was performed with FastTree2.1.10 [52], using the gamma and general time-reversible model parameters. NCBI accession numbers for the reference genomes used in the genealogy are: AE008922.1 (*Xanthomonas campestris*), NC_012660.1 (*P. fluorescens* SBW25), NC_002947.4 (*P. putida* KT2240), NZ_PXJI00000000.1 (*P. otitidis* PAM-1), NC_002516.2 (*P. aeruginosa* PAO1), NZ_FOFP00000000.1 (*P. cuatrocienegasensis*),

### Genomic and functional analysis

In order to study the impact of desiccation on genomic and functional diversity, the global core, cores by sampling, exclusive accessory and shared accessory genomes were obtained with the consensus of three clustering algorithms (BDBH, OMCL and COG) of GET_HOMOLOGUES [53], and aligned with ClustalW [54]. The orthologous genes of the global core genome were used to obtain the nucleotide diversity (π) and average Tajima’s *D* with DnaSP v5 [55]. Functional annotation of the genes of the exclusive accessory genomes was done with Blast2GO [56]. Biological Process gene ontology category was used for this analysis.

### Demographic analysis

Coalescent simulations were carried out and evaluated to find the most probable demographic history of *P. otitidis* from Laguna Intermedia considering the desiccation events. The raw reads were filtered by quality with Sickle v1.33 [57] and mapped against the genome of *P. otitidis* PAM-1 with the Burrows-Wheeler aligner [58].

For the SNP calling, the Genome Analysis Toolkit [59], SAMtools [60] and VarScan2.3.9 [61] programs were used. The folded frequency spectrum was calculated with easySFS (https://github.com/isaacovercast/easySFS).

Coalescence simulations were performed with fastsimcoal2 [62]; 500 repetitions of 500,000 simulations were carried out with 40 optimization cycles and 1000 parametric bootstraps of the best model to obtain 95% confidence intervals. Demographic models allow the possibility of positive (expansion: E) and negative (contraction: K) population changes as well as absence of changes (constant population size: C) in three time intervals (August 2003 to October 2011; October 2011 to November 2012; November 2012 to October 2015) resulting in 27 different models. The initial parameters used in the models were: generational time of *P. aeruginosa* in low phosphorus medium = 0.04 h-1 [63], range of generations from 2003 to 2015 = 6300 to 2100 (this study), mutation rate of *P. aeruginosa* wild type = 5 e-9 [64], recombination rate calculated from R/θ = 2.5 e-8 (this study), effective population size range = 500,000 to 8,000,000 [45]. The evaluation of the models was done by calculating and comparing their weighted AIC values (wAIC) which serve as a measure of the strength of each model in comparison to the others, given their likelihoods, from 0 to 100%.

### Evolutionary rescue

Evolutionary rescue is a phenomenon on which populations contract near to extinction as a result of a strong sudden environmental change, but then recover due to changes in absolute allele frequencies and the appearance of novel selective pressures [16–18]. Positive selection on populations that undergo such demographic contractions followed by an expansion, particularly when a variant increases in frequency in a population, can serve as evidence of the occurrence of this phenomenon. To find signals of positive selection on the core genome of *P. otitidis*, we used the McDonald-Kreitman test [65], built-in in the R package PopGenome v2.7.1 [66] on the population from 2003 against the population from 2015.

### Recombination estimation

To assess the changes in the importance of HGT, recombination rate according to θ (R/θ), insert size (δ), number of substitutions introduced per insert (δν) and the effect of recombination vs mutation on genome level variation (*r/m*) were calculated with ClonalFrameML [67]. 1000 pseudoreplicates were performed to obtain 95% confidence intervals.

## Results

### Genomic variation decreased after the perturbation

The genealogy of core genomes shows that there is phylogenetic correspondence between isolates from 2003 and 2015, which makes the comparison between them valid. In addition, it confirms the proximity of all isolates with *P. otitidis* PAM-1 (Fig 1).

**Figure 1.**
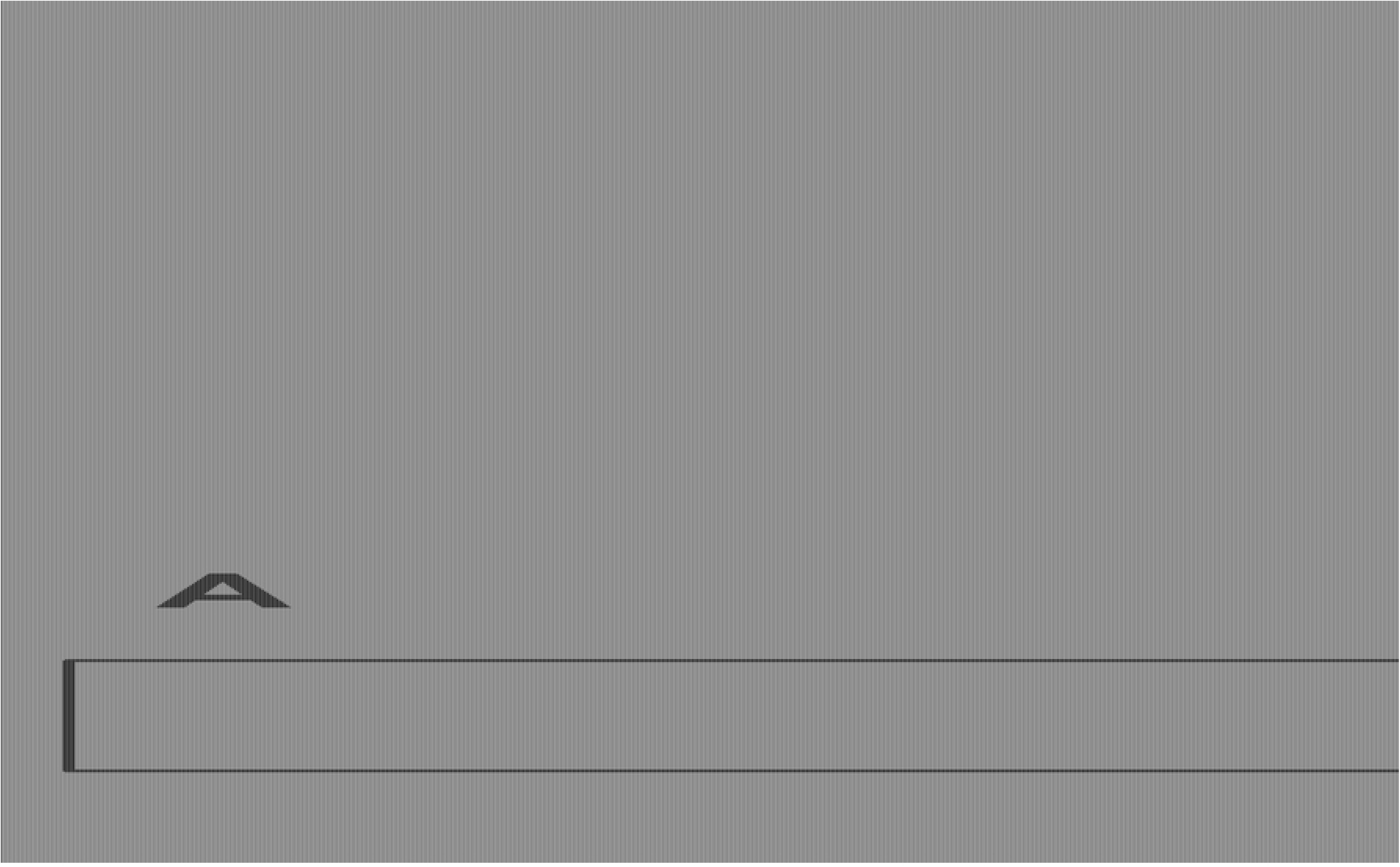
Genealogy of core genomes of *Pseudomonas otitidis* collected from Laguna Intermedia in the Churince system. (A) Genealogy of the Pseudomonas genus including multiple outgroups and (B) zoomed at the clade that comprises the Churince isolates from this study. The isolates from August 2003 (original state) are in blue and from October 2015 (perturbed state) in red. The bar below the trees represents branch lengths. Both phylogenies were calculated by FastTree2.0. All genomes were aligned with progressiveMauve and the core genome was extracted with the stripSubsetsLCBS script (Darling et al, 2010).

Genomic variation of *P. otitidis* from Laguna Intermedia suffered a significant decrease both in its gene and nucleotide diversity (π). The core genome increased by 17.3% (559 genes) while the accessory decreased by 32.8% (486 genes). However, the total gene pool increased by 4.6% (242 genes) (Fig 2), possibly by HGT due to an increase in recombination and SOS response, as discussed further.

**Figure 2.**
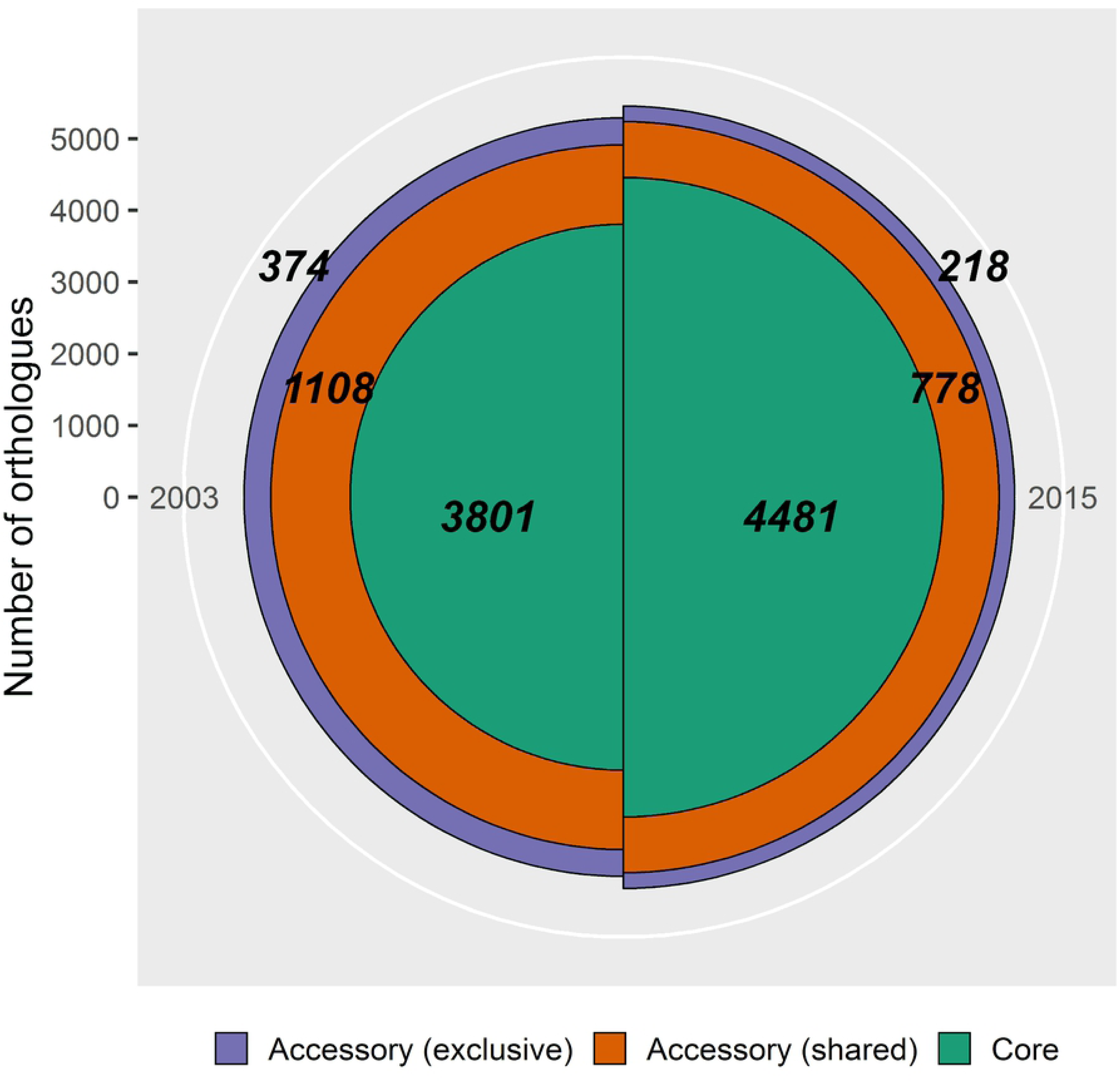
Pan-genomic comparison of *Pseudomonas otitidis* from 2003 and 2015 from Laguna Intermedia in the Churince system. The population from 2003 consists of 5 individuals and has a pan-genome of 5283 genes and the one from 2015 consists of 7 individuals with a pan-genome of 5477 genes. The green fraction represents the core genome, the orange one the shared accessory genome and the purple one the exclusive accessory genome of each sampling. Bold numbers indicate the quantity of genes in each category.

The average π of the global core genome (3682 genes) decreased from 0.0075 to 0.0058, which represents a significant loss of 22% of its nucleotide diversity (paired two-sided Wilcoxon test: V = 4.42e6, p <0.001), as well as a significant decrease in its variance from 1.39e-4 to 6.4e-5 (Brown-Forsythe test: F (1, 7362) = 27.6, p <0.001) (Fig 3).

**Figure 3.**
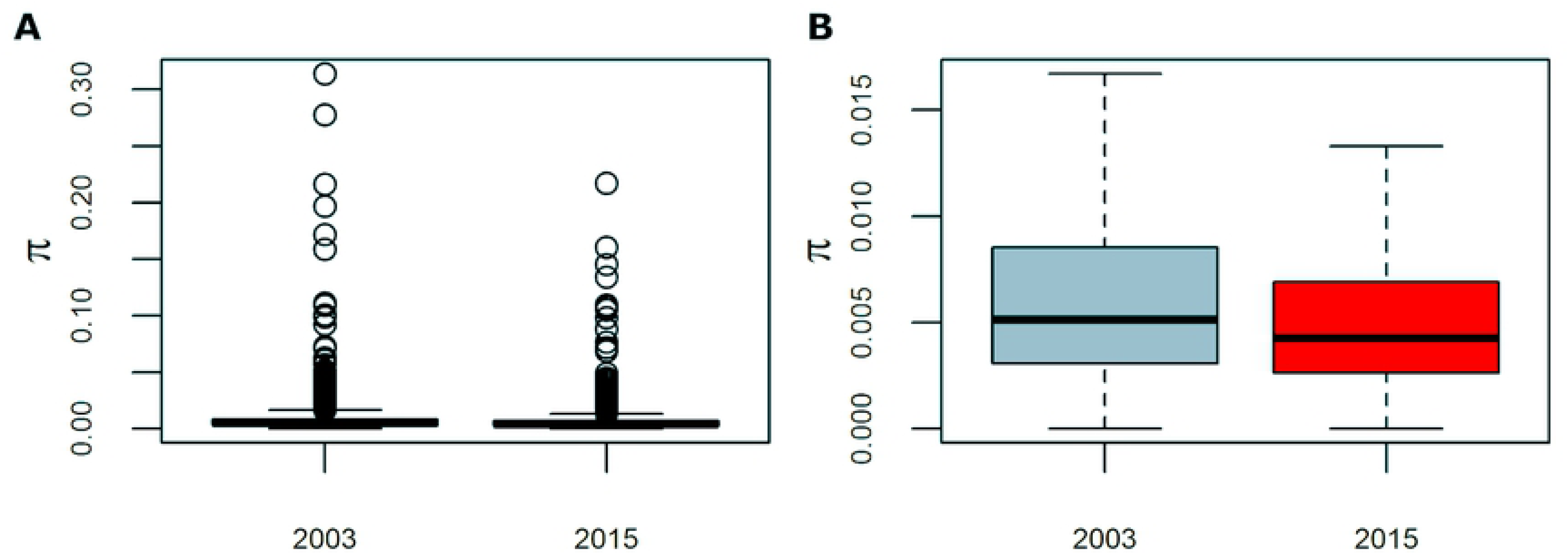
Barplots of the average nucleotide diversity (π) of *Pseudmonas otitidis* at the global core genome level. The global core genome consists of 3682 genes. (A) Barplots presented without and (B) with outliers (values outside 1.5 times the interquartile range).The change of mean and variance of π are significant (paired two-sided Wilcoxon test: V = 4.42e6, p <0.001; Brown-Forsythe test: F (1, 7362) = 27.6, p <0.001).

### The demographic history of a lineage bloom and subsequent demise

The two best coalescent models according to their wAIC values are contraction-expansion-contraction (KEK, wAIC = 73.15%) and contraction-expansion-constant (KEC, wAIC = 18.43%), and both coincide in an initial population contraction from 2003 to the first desiccation event in 2011 of more than 99%, followed by a drastic population increase until the second desiccation event in 2012 (Fig 4, S1 Table). The 95% CI overlap in both models reach a similar final population size in 2015, which is approximately 1000 times larger than its initial size.

**Figure 4.**
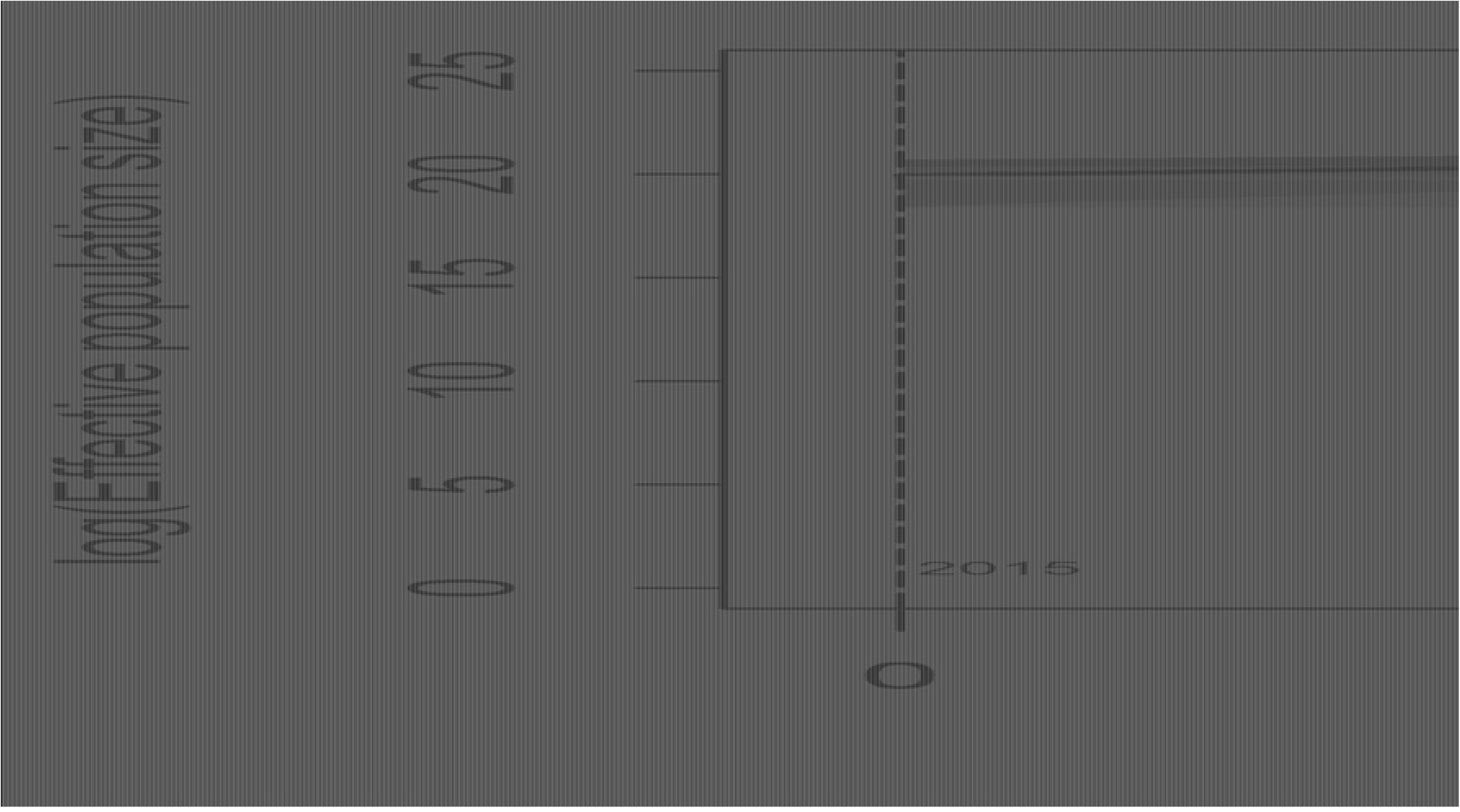
Graphical representation of the best two demographic models according to their weighted AIC value based on the results obtained from fastsimcoal2. Coalescent simulations were performed using the observed allele frequency spectrum of the *Pseudomonas otitidis* population from 2015, taking into account an initial population range of 500,000 - 8,000,000 estimated by Avitia et al. [45], mutation rate of wild type *P. aeruginosa* of 5e-9 calculated by Oliver et al. [64], generational time of *P. aeruginosa* in low phosphorus medium of 0.04 h-1 calculated by Buch et al. [63], and range of generations from 2003 to 2015 of 6300 to 2100 and recombination rate (R) from R/θ = 2.5 e-8 calculated in this study. The solid colored lines are the average values and the shaded areas represent the 95% confidence intervals. Contraction-Expansion-Contraction (KEK) is the best model with a wAIC value of 73.15% (green) and Contraction-Expansion-Constant (KEC) follows with 18.45% (orange). Dotted vertical lines mark the desiccation events according to the generations calculated by fastsimcoal2. The population contraction from 2003 to 2011 for both models was approximately 99%. For detailed values see Supplementary Table 1.

There was a significant increase in the average Tajima’s *D*, which went from a marginally negative value (*D* = −0.03) in 2003 to a positive value (*D* = 0.26) in 2015 (paired two-sided Wilcoxon test: V = 2.07e6, p < 0.001). Variance in *D* increased from 0.50 to 0.61, and the change was also significant (Lavene’s test: F (1,7004) = 23.8, p < 0.001) (Fig 5). After the sampling of 2015, no other pseudomonad was recovered from the lagoon, even if it was sampled at least two times per year every year until it dried two years later.

**Figure 5.**
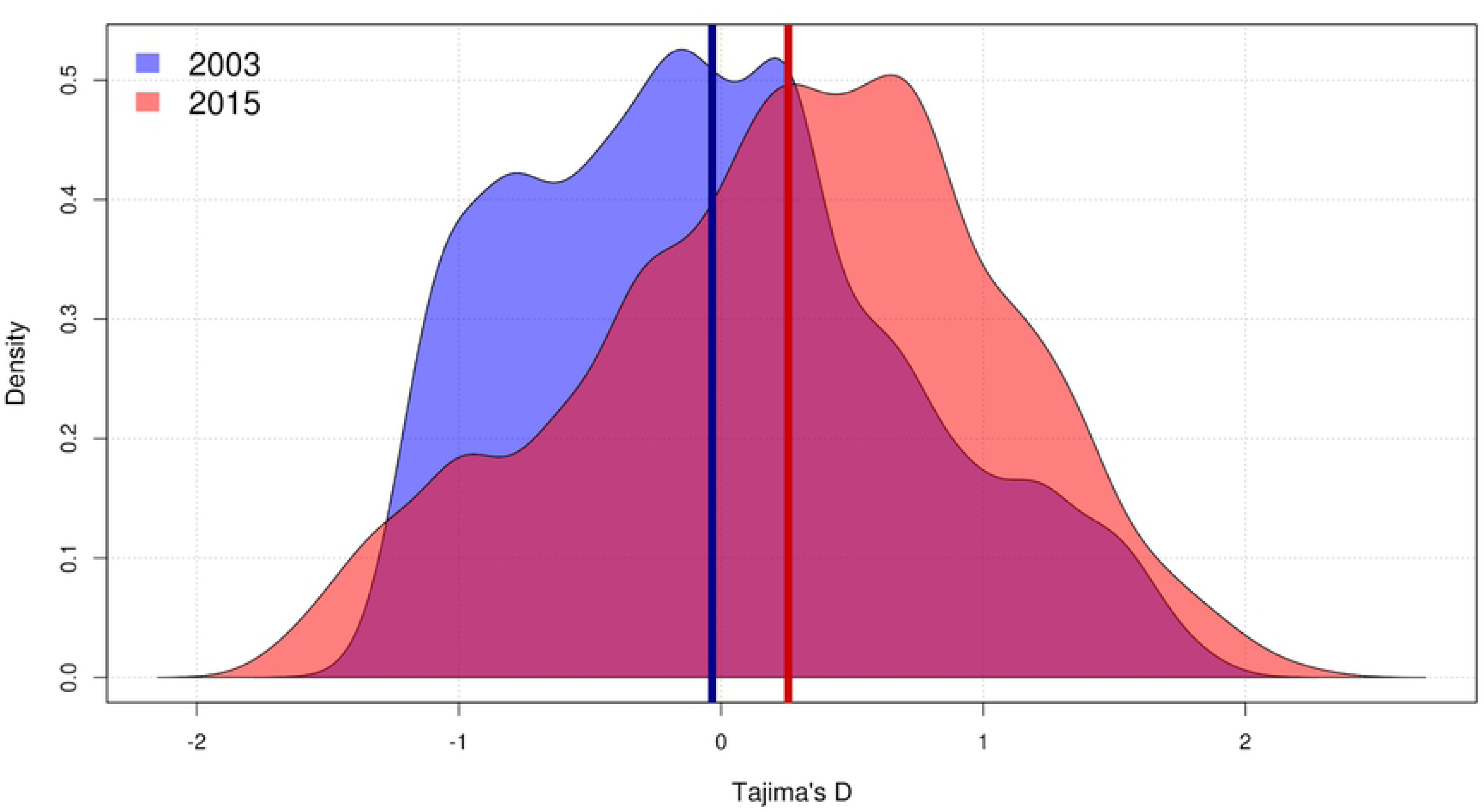
Density plot of the distribution of *Pseudomonas otitidis* Tajima’s *D* at core genome level. Populations from 2003 and 2015 consist of 5 and 7 individuals, respectively. Only 3503 genes were used for the analysis as the rest of the 3682 global core doesn’t have any variation to measure. Kernel smoothing was used to plot the values. Vertical bars represent the means: −0.03 in 2003 (blue) and 0.26 in 2015 (red). The decrease in mean value and the increase in variance of are significant (paired two-sided Wilcoxon test: V = 2.07e6, p <0.001; Lavene’s test: F (1,7004) = 23.8, p <0.001).

### Evidences of evolutionary rescue: positive selection

A McDonald-Kreitman test was performed on all 3682 genes of the global core genome between the population of 2003 and 2015. Positive selection was detected only on amino acid 36 (Neutrality Index = 0) of the C4-dicarboxylate transport sensor protein DctB, which senses carbon sources such as succinate, malate and fumarate, and interacts directly with the DctA protein which transports these compounds into the cell [68]. On a population level, the codon of this amino acid coded for arginine and histidine in 2003 but only for arginine in 2015. All genetic variation at that specific codon was lost, suggesting the bloom of a successful clone.

### The *P. otitidis* functional shift in a perturbed environment

The exclusive accessory genome comprises the unique genes of each sampling, those that were lost (only found in 2003) and gained (only found in 2015), with a net decrease of 41.7% (156 genes). Exclusive accessory genomes of both samplings (Fig 2, purple fraction) were functionally annotated (Tables S2 and S3), with 27.8% of genes (104 genes) for the population of 2003 and 36.7% (80 genes) for the population of 2015 being functionally identifiable under the Biological Process gene ontology category. According to a multi-level functional analysis performed by Blast2GO, included functions related to metabolism of organic nitrogen compounds and small molecules, modification of macromolecules, phosphorylation and recombination. In contrast, genes involved in regulation of cellular process, biosynthesis of cellular macromolecules and nucleobase-containing compounds and DNA and cellular metabolic process were lost (Fig 6). Loss of biosynthetic and regulatory genes has been associated with bacterial adaptation to a new environment [69–70] while phosphorylation-related functions and DNA recombination have been observed to function as an stress response against environmental stress [71–73]. A larger number of small-molecule metabolic genes is correlated with an increase in metabolic versatility in a nutrient-diverse environment and a free-living lifestyle [74]. Even though all other functional categories decreased in their number of genes, genes with transport functions, particularly of organic substrates, increased by 42.8%. (Fig 6), suggesting its possible adaptive relevance function in an environment with higher nutrient availability [75].

**Figure 6.**
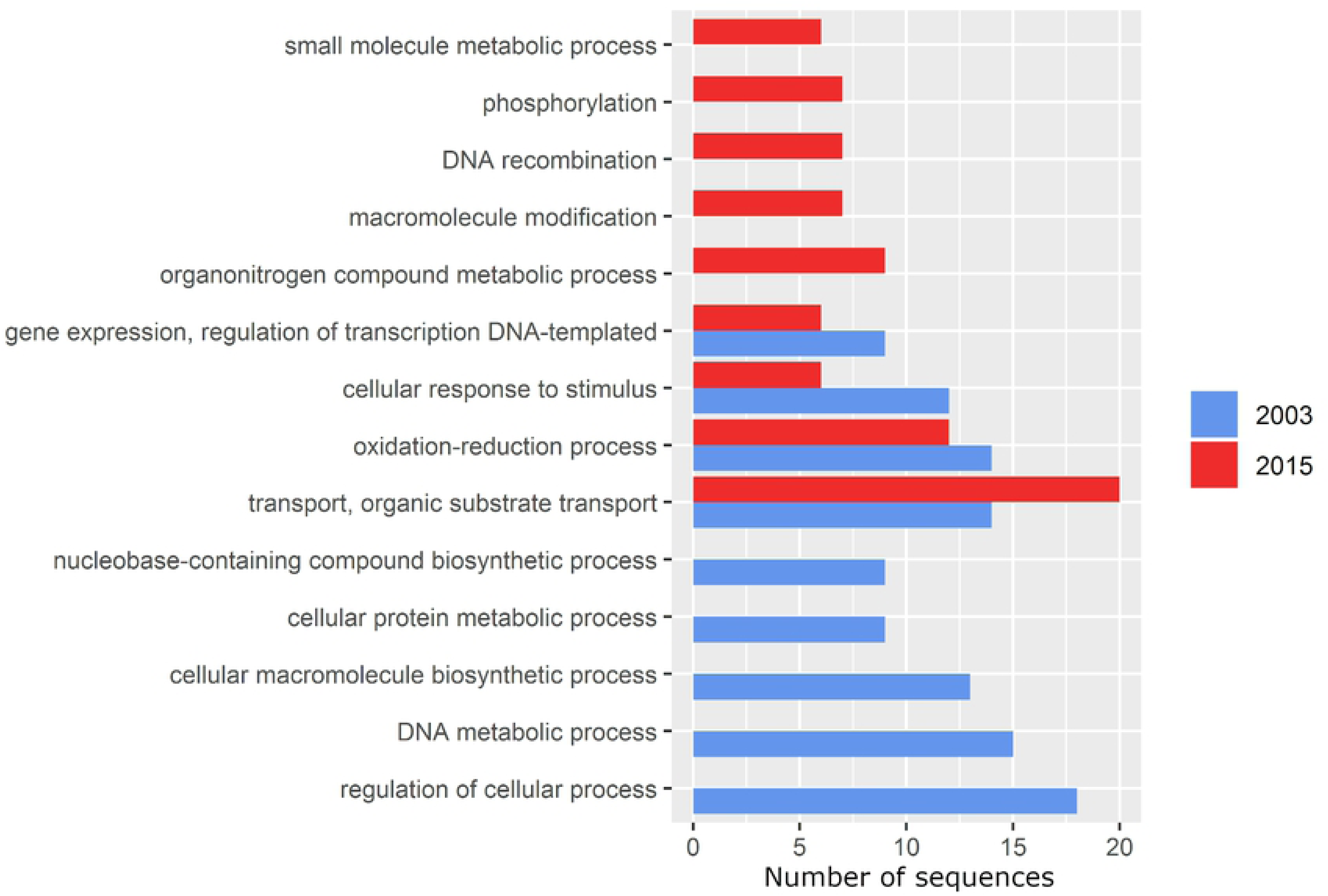
Multi-level gene ontology graph of the exclusive accessory genome of *P. otitidis* under the Biological Process category calculated by Blast2GO. The gene ontology terms (functions) are on the Y axis and the X axis represents the number of sequences assigned to each one. The 2003 and 2015 populations are marked in blue and red, respectively. Bars alone represent functions lost (2003, blue) and gained (2015, red).

### Recombination increased in a more homogeneous *P. otitidis* population

As could be expected from the gain of recombination related genes, the genomic diversity explained by recombination (R/θ) increased slightly but significantly (0.19, CI 95% = 0.00017 to 0.20, CI 95% = 0.00018) while the size of the insert decreased significantly in a 23.6% (110 pb, CI 95% = 0.085 to 84 pb, CI 95% = 0.063), as well as the number of substitutions per insert that decreased significantly in a 19.2% (5.5, CI 95% = 0.0043 to 4.5, CI 95% = 0.0035). As a result, the ratio of recombination and mutation (*r/m*) also decreased significantly in a 12.4% (1.05, CI 95% = 0.0011 to 0.92, CI 95% = 0.0009) as the population became more homogeneous.

## Discussion

### The loss of genetic diversity and function as consequences of habitat loss

Environmental perturbation due to the overexploitation of the deep aquifer at CCB has a direct effect not only on aquatic macroorganisms, such as turtles and fishes [76, 77], but also on microbes, as reported previously in studies of community interactions [25, 35]. In the present study we documented genomic and demographic changes in the population of environmental *P. otitidis* as well as its evolutionary response to three desiccation events.

Although all *P. otitidis* isolates are close members from the same lineage (Fig 1), the population sample from 2015 had a larger genomic core and smaller accessory genome (Fig 2), alongside a significantly lower average and variance of π than the sampled strains from 2003 (Fig 3). This generalized decrease in genomic variation along with the loss and gain of some gene functions could be expected given the ∼99% population bottleneck of 2011 followed by the approximately 1000 fold population expansion of 2015 estimated by our coalescent models (Fig 4, S1 Table). Population bottlenecks decrease genetic variation by the stochastic elimination of low frequency alleles, which impacts negatively on the rate of accumulation of adaptive variation [78–80]. Moreover, if the length of a bottleneck is protracted, population size will decline and may reach extinction, depending on its growth rate [18]. Although *P. otitidis* managed to escape the bottleneck of 2011 and was thriving during the second and third desiccation events of 2013 and 2015, the lineage was never sampled again after October 2015, even if we intensely looked for it, possibly due to its extinction.

The first bacteria that apparently became locally extinct in the Churince system was *Bacillus coahuilensis* [30, 81], a planktonic marine bacillus from Laguna Grande that has not been found after 2006. However, in that case, the outcome could be easier to predict, given the possible black queen dynamics [82] in *B. coahuilensis,* which was a specialized lineage with a particularly small genome, lacking several metabolic pathways, therefore, an organism highly dependent on several metabolites from other community members [30,81,83]. What was observed in our study is more intriguing, since *P. otitidis* is a planktonic generalist that was observed consistently over the years at Churince [43–45, 33, 35, 36] and even experimented a dramatic population increase just before apparently going locally extinct.

### How to survive extreme stress, or at least buy some time

The story of this environmental drama is told by the estimated changes in population sizes between 2003-2015. According to the two best demographic models (KEK, wAIC = 73.15% and KEC, wAIC =18.43%), we observed a population bottleneck which drove *P. otitidis* near to extinction in 2011, followed by a drastic population expansion that peaked in 2013 (Fig 4, S1 Table). Together with the significant decrease in genomic variation and the significant increase in Tajima’s *D* and its variance [84, 85], the evidence strongly suggests the occurrence of an evolutionary rescue event. Therefore, we believe that evolutionary rescue changed *P. otitidis* niche, from a species adapted to low nutrient conditions to a scavenger specialized strain. Evolutionary rescue has been suggested to occur in microbial populations under a sudden environmental change, and in particular has been documented in *P. fluorescens* subjected to streptomycin, where an sharp initial population decrease was observed before the rapid evolution of resistance [86]. In those cases, the selective pressure is severe enough to lower the population’s mean fitness below one, causing the population size to decrease near extinction. However, these populations recover as a beneficial allele increases its absolute frequency, helping them to overcome and adapt to the new environmental conditions [17].

The McDonald-Kreitman test [65] found a single positive selection signal at amino acid 36 of the DctB protein (Neutrality Index = 0). While Eyre-Walker [87] points out the risks of the literal interpretation of the test result when a sudden population expansion occurs given that slightly deleterious mutations that currently do not segregate in the population and may have been fixed in the past can show as artifactual evidence of adaptive evolution, in this study, this particular positive selection signal is consistent with the change of niche from a strain that was efficient in an oligotrophic environment and that perhaps was even a community cooperator, to a scavenger in a nutrient-rich environment. This eco-evolutionary change is also supported by the acquisition of genes with transport functions particularly of organic substrates [75], on the exclusive accessory genome of the surviving lineage (Fig 6). The *dctB* gene under positive selection codifies for a protein that is a transmembrane sensor of C4-dicarboxylate carbon sources, such as succinate, malate and fumarate that interacts with the DctA transporter protein, for the internalization of these molecules to the cell [68], and seem to be a strategic gene for surviving given the increase of availability and concentration of nutrients in Laguna Intermedia, as during the desiccation many animals died, and this new source of nutrients would mean the opportunity of local adaptation for a scavenger *P. otitidis*. Selective sweeps [19] are part of an evolutionary rescue scenario [17] and this seems likely given the complete loss of variation on the codon of this particular positively selected amino acid.

The role of adaptive gene loss in bacteria has been previously studied [88]. In this study, genes with functions such as biosynthesis and regulation were lost in the *P. otitidis* population after the bottleneck (Fig 6). Loss of biosynthetic genes in bacteria is a general and well documented phenomenon by which lineages become auxotrophic in particular environmental conditions, apparently due to the cost of maintaining those genes, or because the genes accumulate mutations if they are not used, and thus become unable to produce some of metabolites they need to grow in most conditions [89]. This may be an adaptive response when the corresponding metabolite is sufficiently present in the environment [82]. In this particular case, the sudden overabundance of nutrients and cellular compounds in the remaining water was caused by dead fishes and other animals that could not survive the desiccation events. We also believe the disruption of the original community interactions and their microbial “market”, may have rendered biosynthesis of certain metabolites an unnecessary burden, thus being lost as an adaptive response [69]. Albalat and Cañestro [70] pointed out the importance of loss of regulatory functions in the adaptation of bacterial populations to new environments that require alternative or new metabolic pathways, which seems to be the case at Laguna Intermedia given the severity of the perturbation.

Functions gained include DNA recombination and phosporylation, both of which have been observed to play a role in environmental stress response. The gain of genes involved in DNA recombination after the selective sweep is interesting, and correlate with a significant increase in recombination rate (R/θ, 0.19 ± 0.00017 to 0.20 ± 0.00018, p <0.001), pointing to a possible mechanism of evolutionary rescue by HGT. Bacteria such as *Escherichia coli* [90], *Vibrio* [15], *Staphylococcus* [91] and also *Pseudomonas* [92] have a stress response known as SOS that consists on repairing DNA damage caused by multiple sources of stress such as UV light, fungal metabolites, reactive oxygen species [72], exposure to antibiotics and phages [93]. SOS response can facilitate evolutionary rescue by integrating exogenous DNA (that arrived through HGT) to their genome. We believe that the multiple desiccation events on Laguna Intermedia likely triggered a SOS response in *P. otitidis*. This in turn could explain the increase in the size of the gene pool in 2015 due to the incorporation of exogenous DNA. However, *r/m* decreased significantly, suggesting that although recombination increased, it occurred between a less diverse gene pool, which coincides with the loss of diversity observed and that closer bacterial lineages tend to recombine more frequently [94, 95]. Phosphorylation is a key mechanism involved in signal transduction as a response to environmental stimuli such as nitrogen deprivation, phosphorous availability, osmolarity and chemotaxis molecules [71]. Moreover, experimental studies in rhizobia indicate that phosphorylation plays an essential role in many physiological processes in adaptation to various environmental factors (e.g., exopolysaccharide and flagellum production, dicarboxylate transport, catabolite repression, phosphate utilization, N2 fixation, and adaptation to pH stress and microaerobic conditions) [96] through multiple regulatory mechanisms such as phosphoenolopyruvate-dependent phosphotranspherase systems, Hanks-type serine/threonine kinases and serine/threonine phosphatases [73]. As new stimuli surged in Laguna Intermedia, the gain of phosphorylation-related genes may have helped *P. otitidis* to withstand the new environmental conditions and adapt to them. Additionaly, the increase in small-molecule metabolism genes correlates with an increase in metabolic versatility required under conditions of high competition [97], and a large variety of nutrients [74], supporting the hypothesis of the lifestyle change of *P. otitidis* to a scavenger generalist.

## Conclusion

### The tale of loss

From an evolutionary rescue event to the extinction of a bacterial lineage, this study provides evidence on the eco-evolutionary consequences of anthropogenic perturbation over the environmental microbiota. *P. otitidis* from Laguna Intermedia went through a drastic population bottleneck that preceded a population expansion of 1000 times its original size in 4 years. This perturbation was a result of the overexploitation of water for alfalfa irrigation and human consumption. Also, its diversity (nucleotidic and pan-genomic) decreased while Tajima’s *D* increased in average value and variance. The finding of an amino acid under positive selection of a protein involved in the sensing and transport of carbon sources, alongside the number increase of transport genes in its exclusive accessory genome, suggests that the occurrence of a selective sweep due to an instance of evolutionary rescue in response to the desiccation events. In addition, during the perturbations, recombination rates increased and genes with recombination-related functions were gained, suggesting an increase in the activity of SOS responses. Lastly, the adaptive loss of biosynthetic and regulatory genes as well as the acquisition of phosphorylation and small-molecule metabolism genes supports the hypothesis of niche change from a community cooperator to a scavenger lifestyle. However, this rescue left genetic diversity very low, and the specialist scavenger could not adapt to the next phase, in a mostly dry sediment cracked lagoon.

Efforts to save CCB wetlands are in effect right now. Maybe there is hope that the Churince system may resurge from those dry sediments. Dramatic changes in water usage in arid lands are needed and this wetland may shows us how to change policy through science.

## Acknowledgements

Manuel II García Ulloa is a doctoral student from the Programa de Doctorado en Ciencias Biomédicas, Universidad Nacional Autónoma de México (UNAM), and has received CONACyT fellowship 596613. We greatly acknowledge V.M.D Gabriel Manuel Rosas Barrera and Dr. Gabriel Yaxal Ponce-Soto for the help during the sample collection and lab assistance. We also thank Dra. Erika Aguirre Planter and Dra. Laura Espinosa Asuar for the technical support.

## Supporting information

**S1 Fig. Barplots of environmental measurements from Laguna Intermedia since 2002 to 2016.** Temperature is in °C (A), salinity in ppt (B), pH (C), dissolved oxygen in mg/L (D) and specific conductivity mS (D). Measurements before 2015 were taken by Escalante et al. [43], Ponce-Soto et al. [36] and García-Ulloa et al. [36]. Desiccation events are indicated by ***.

**S1 Table. Values for parameters of the two best demographic models calculated by fastsimcoal2**. Number of generations, effective population size and 95% confidence intervales for each one were estimated for the initial *Pseudomonas otitidis* population (2003) and on each desiccation event (2011, 2013 and 2015), for the Contraction-Expansion-Contraction (KEK) and Contraction-Expansion-Constant (KEC) models.

**S2 Table. Functional annotation of the exclusive accessory genome of the *Pseudomonas otitidis* population from 2003.** Annotation by gene ontology was done with Blast2GO. 184 (49%) of 374 of sequences were annotated.

**S3 Table. Functional annotation of the exclusive accessory genome of the *Pseudomonas otitidis* population from 2015.** Annotation by gene ontology was done with Blast2GO. 135 (62%) of 218 of sequences were annotated.

